# Effect of temperature and loading frequency on the performance of vertical subsurface flow constructed wetlands: Modelling study using HYDRUS

**DOI:** 10.1101/2021.03.25.436912

**Authors:** Nikita Jayswal, Jorge Rodríguez

## Abstract

Constructed wetlands are used in wastewater treatment for the removal of organic matter, nutrients, pathogens and other pollutants from water. The diversity of chemical, physical and biological processes taking place inside constructed wetlands (CWs) make them difficult systems for accurate quantitative description through conventional modelling approaches. However, existing modelling tools can bring valuable insight into important phenomena and interactions that might remain otherwise neglected. In this study the CW2D module under HYDRUS – 2D is used to identify the possible mechanisms by which temperature and loading frequency impacts the performance of the vertical subsurface flow constructed wetlands (VSSFCWs). An improvement on CWs design and operation guidelines in warm regions is intended. Case studies of a VSSFCW at 10°C, 20°C and 30°C treating septic tank effluent are simulated with parameters adjusted for high temperature conditions. The simulation results suggest oxygen limitation as possibly constraining nitrification capacity at high temperatures. Simulations also suggest that higher loading frequencies enhance performance and capacity. This detailed mechanistic modelling study aims at bringing insight into possible non trivial interactions and mechanisms in CWs such that in conjunction with a necessary experimental validation, should lead to enhanced operation and design in warm regions.

## Introduction

Constructed wetlands (CWs) are widely used to treat wastewater and improve water quality. Their mixture of water, plant roots, litter and a variety of microorganisms provide good conditions for the removal of organic matter, nutrients and decreased concentration of toxic trace metals, organic chemicals and pathogens (Langergraber and Šimůnek, 2005). The multiple physical, chemical and biological processes operating simultaneously, in interaction with each other, make CW systems complex and difficult to model. As a consequence, CWs are often approached as “black boxes” used for the purification of water (Langergraber *et al*., 2007). CW designs had for long been based on rule of thumb approaches using specific surface area requirements (Brix and Arias, 2005) or first order decay models (Kadlec *et al*., 2000). Numerical models have been used mainly to provide some insight into the processes taking place inside the system.

Vertical subsurface flow constructed wetlands (VSSFCWs) have been demonstrated to achieve high ammonia nitrogen removal efficiencies when loaded intermittently (Langergraber, 2008) with water infiltrating into the substrate and draining down vertically where it is collected in the drainage network (Langergraber and Haberl, 2001; Pucher and Langergraber, 2018). High rate of oxygen transfer is achieved as air enters at every loading making this intermittent operation suitable for nitrification and aerobic organic matter (OM) removal processes.

Temperature is a key factor on the rate of the microbially mediated reactions involved in wastewater treatment and therefore on the performance of biological treatment processes, including CWs, which therefore can exhibit large seasonal variability. The optimum temperatures for treatment performance are known to depend on multiple interacting variables with higher temperatures increasing all (microbially mediated) reaction rates. Temperatures above 15°C have been reported to enhance organic matter (OM) removal efficiencies in CWs as well as the microbial activity responsible for the removal of COD, BOD, TKN, NH_3_–N and phosphorus in horizontal SSFCW (Akratos and Tsihrintzis, 2007). High temperatures however decrease oxygen solubility. Dissolved oxygen (DO) levels are critical for the aerobic microbial activity responsible for OM removal and nitrification. DO limitation for OM removal in CWs has been widely reported (Chang *et al*., 2012; Hiley, 1995; Vymazal and Kröpfelová, 2009). In the case of nitrification it has been reported to be inhibited at temperatures below 10°C due to the decline in the growth rate of nitrifying bacteria (Werker *et al*., 2002). However at the same time nitrification correlates with DO concentrations in the wetland and therefore might become also limited at high temperatures (Chang *et al*., 2012). Other microbial activities such as anoxic denitrification, part of the complete CWs nitrogen removal cycle, have also optimum performance temperature ranges, reported between 35°C and 45°C (Kadlec and Reddy, 2001).

Most experimental studies on the effect of temperature or seasonal variation on the treatment performance of CWs describe the effect of low temperature variations on effluent quality comprehensively (van de Moortel *et al*., 2010; Akratos and Tsihrintzis, 2007; Stefanakis and Tsihrintzis, 2012; Kadlec and Reddy, 2001) with generally less emphasis on the high temperature scenarios.

In this work a model simulation-based study, using the advanced numerical model HYDRUS CW2D, is conducted to evaluate the impact of high temperature on the performance of CWs. The study is applied to a model single stage VSSFCW treating typical post septic tank composition effluent. Simulations are conducted at three different temperatures namely 10, 20 and 30°C. The aim is to understand the temperature effect on the treatment performance (mainly in terms of effluent quality as COD, NH_4_–N) and the likely associated mechanisms as well as to provide suggestions for the optimum operation in warm climates. The investigation of the temperature-related mechanisms and the operation optimisation are made only on the basis of the simulations results and will require experimental verification.

## Materials and Methods

The biokinetic CW2D wetland module of HYDRUS was used in this study. Details of the CWs system model implementation and parameters are explained in detail below.

### Model CW system

A one-stage VSSFCW system was selected for the study. The geometry implemented in HYDRUS consisted of a basin of 1 m^2^ area with a filter (main) layer of 35 cm and a drainage layer of 15 cm in depth (see Supplementary Figure S1 for detail). The influent to the CW is defined as from representative domestic wastewater after typical septic tank reductions of only OM and solids (Abdel-Shafy and El-Khateeb, 2013), septic tanks typically leave the nitrogen loads unchanged. Table 1 shows the assumed model raw wastewater and the CW influent composition after typical assumed septic tank pre-treatment. The influent composition is expressed in terms of the reactive transport components of the HYDRUS CW2D module (see Table 1). The COD fractionation in terms of CW2D components was conducted based on (Pasztor et al., 2008).

**Table 1.**
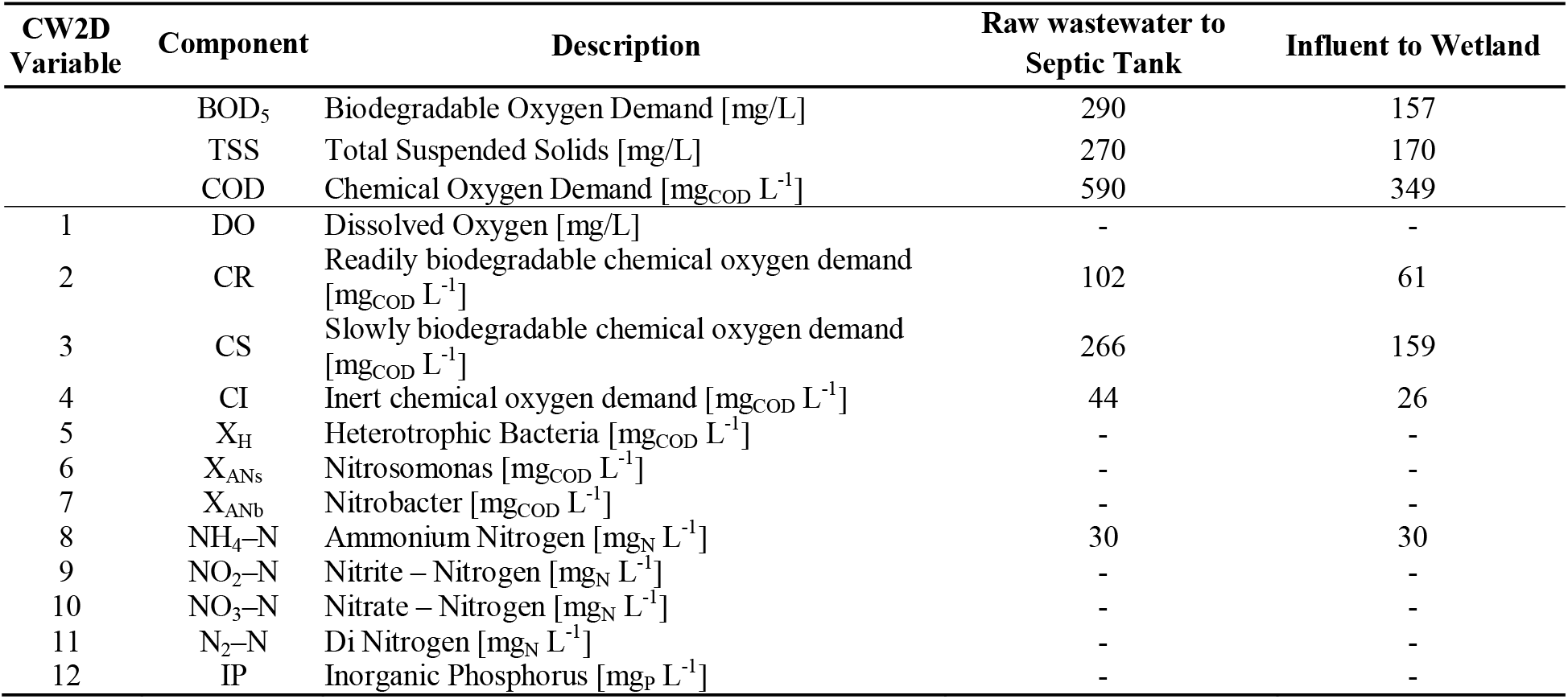
Selected model wastewater characteristics (Metito, 2013) in terms of CW2D model components

Population Equivalent (PE) units based on 60 g BOD/person/day are used (Henze, 2008). A CW hydraulic loading rate of 0.2 m^3^/m^2^/day with a BOD concentration of 157 mg/L was used as reference load value. This is equivalent to an area of 2 m^2^/PE approximately, value within the conventional load ranges of septic tank integrated CWs (EPA Ireland, 2009).

### CW2D biokinetic model overview

The multi-component biokinetic module CW2D of HYDRUS is based on the Activated Sludge Models (ASMs) (Henze *et al*., 2006). CW2D can model the biochemical transformation and degradation processes for organic matter, nitrogen and phosphorus (Langergraber and Šimůnek, 2005).

The CW2D module incorporates twelve reactive transport components (see Table 1) and nine reaction processes including hydrolysis, mineralization of OM, nitrification as a two-step process, denitrification and lysis for microorganisms. All the OM is assumed to be in aqueous phase and all reactions are also assumed to take place in the aqueous phase. The CW2D also assumes the microbial biomass to be immobile and present only in the solid phase with the microbial decay and sink processes lumped together under lysis.

All biochemical transformation and degradation processes are modelled in the CW2D using the Monod rate equations as defined in ASM models. The growth and decay rate parameters as well as the diffusion coefficients are temperature dependent and are estimated. In the study, kinetic parameters for 10°C and 20°C from ASM models are used (Henze *et al*., 2006) while those for 30°C are estimated as per Tchobanoglous *et al*. (2003). The saturation/inhibition coefficients are considered constant for all temperatures. The coefficient for hydrolysis at 30 °C is estimated using the guidelines from (Langergraber, 2007). Additional information regarding the parameters used can be found in the supplementary material (Table S1). A comprehensive description of the CW2D model can be found in (Langergraber and Šimůnek, 2005).

### HYDRUS model description

The HYDRUS software package solves Richard’s equation for unsaturated – saturated water flows and convection – dispersion equation for solute and heat transport (Langergraber, 2008). There are various models available in HYDRUS for the soil hydraulic properties as well as a variety of system dependent as well as independent boundary conditions.

A 2D geometry in the XZ coordinate is defined for the CW using the HYDRUS interface. Only the main layer of the wetland is considered. The simulation is run for 24 hours and then the results from the last time step are used as the initial conditions to run the model to reach a pseudo steady state. The single porosity van Genuchten–Mualem hydraulic model is used.

The wetland design characteristics are taken from a real two-stage VSSFCW plant located in the UAE Western region. 35cm deep double washed wadi sand with a porosity of 0.3 with a K_s_ of 117 cm/hour in the filter layer is used. The real CW has a 15 cm depth gravel drainage layer with a porosity of 0.41 and K_s_ of 3000 cm/hour, however the drainage layer is excluded from the modelled system. Soil hydraulic parameters from the van Genuchten–Mualem model (Langergraber and Šimůnek, 2005) are used.

A reference wetland load frequency of 8 times per day is used with each feeding loading lasting for 15 minutes. The actual loading rate (precipitation in HYDRUS) is of 10 cm/hour calculated from the number of loadings and the loading time.

Atmospheric boundary condition (time dependent) is applied at the top where the water flows into the wetland. Relative humidity data is required to calculate the evaporation rates and is used from a weather station in Rotterdam for 10°C (Royal Netherlands Meteorological Institute, 2013) and from the Masdar City (Abu Dhabi, UAE) weather station for 20°C and 30°C. HYDRUS atmospheric boundary conditions are used such that solutes leaving the domain from the top boundary do so only by gas diffusion. Free drainage boundary condition is applied at the bottom of the main layer of the transport domain. HYDRUS Type III boundary condition is applied for solute and water transport for both the boundaries.

The initial values of the components used for the simulation are taken as the influent as shown in Table 1. The entire wetland model domain is initialized with these values at time t=0. The influent septic tank pre-treated wastewater is fed to the wetland intermittently (every 3 hours in the reference load frequency). More details on the HYDRUS model set up can be found in Figure S1, Tables S2, S3 and S4 of the supplementary material.

To assess both the impact temperature has on treatment performance indicators (effluent quality) and the possible associated mechanisms, simulations were conducted at three different temperatures. The hydraulic loading of the CW was also changed to push the operation to the limits and obtain insight into the possible underlying mechanisms linked to temperature. Twice (1 m^2^/PE) and half (4 m^2^/PE) the area specific loading rate with respect to the reference value (2 m^2^/PE) were also simulated. All nine cases (three loadings at three temperatures) were simulated for a ten days period and the averaged effluent values from the last simulation day are taken as representative of the pseudo stationary state treatment performance.

## Results and Discussions

Table 2 presents the simulated effluent quality outputs of the CW at different temperatures and loading intensities.

**Table 2.** Simulated effluent characteristics after ten days simulation at different temperatures and loading rates

|  | Temperature | Hydraulic Loading Rate | Area Specific Loading Rate | CR (Readily Biodegr.) | CS (Slowly Biodegr.) | COD | NH<sub>4</sub>-N | NO<sub>3</sub> - N |
| --- | --- | --- | --- | --- | --- | --- | --- | --- |
|  | (°C) | (L/m<sup>2</sup>-day) | (m<sup>2</sup>/PE) | (mg<sub>BOD</sub>/L) | (mg<sub>BOD</sub>/L) | (mg/L) | (mg/L) | (mg/L) |
| **Influent** | - | - | - | 60.90 | 159.38 | 246 | 30.00 | 0.00 |
| **Effluent** | 10 | 100 | 4 | 0.142 | 155.34 | 191 | 0.278 | 38.222 |
| **Effluent** | 20 | 100 | 4 | 0.481 | 19.502 | 61 | 0.017 | 38.075 |
| **Effluent** | 30 | 100 | 4 | 1.279 | 0.075 | 42 | 0.006 | 31.262 |
| **Effluent** | 10 | 200 | 2 | 0.134 | 158.937 | 189 | 1.163 | 31.491 |
| **Effluent** | 20 | 200 | 2 | 0.433 | 40.137 | 75 | 0.027 | 31.271 |
| **Effluent** | 30 | 200 | 2 | 1.094 | 0.811 | 39 | 0.014 | 26.348 |
| **Effluent** | 10 | 400 | 1 | 0.131 | 161.187 | 190 | 11.034 | 19.549 |
| **Effluent** | 20 | 400 | 1 | 0.980 | 76.457 | 108 | 3.424 | 23.955 |
| **Effluent** | 30 | 400 | 1 | 6.902 | 8.733 | 49 | 4.640 | 20.043 |

### Effect of temperature at different loading rates

Figure 1 presents the resulting simulated pseudo stationary state effluent concentrations in terms of readily (CR) and slowly (CS) biodegradable COD as well as ammoniacal nitrogen (NH4-N) for the three loading rates and temperatures.

**Figure 1.**
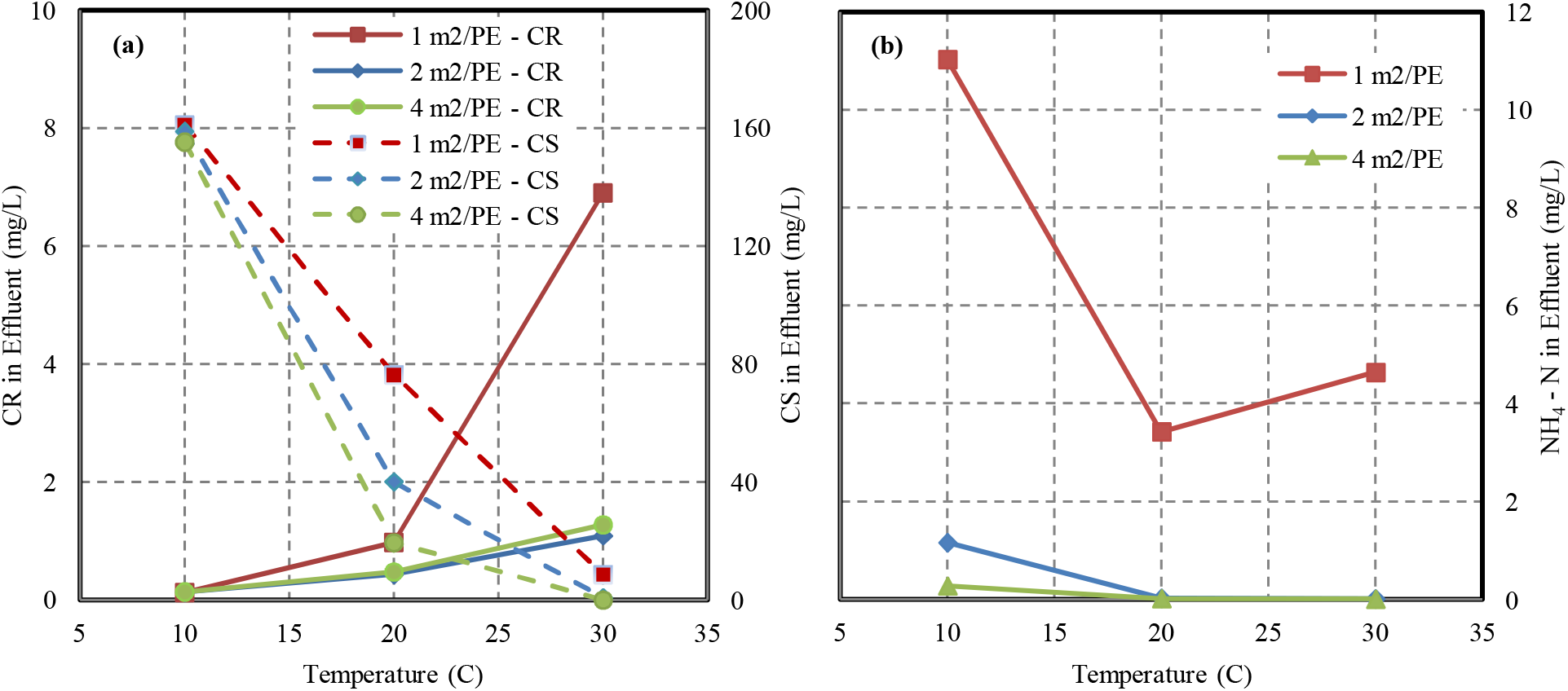
Simulated effect of temperature on the pseudo steady state effluent concentrations of (a) readily (CR) and slowly (CS) biodegradable COD and (b) ammoniacal nitrogen (NH_4_–N) at different specific loading rates.

At the reference loading rate of 2m^2^/PE simulation predict a removal efficiency decrease of both COD (Figure 1.a) and NH_4_–N (Figure 1.b) in the effluent as the temperature increases. Simulations also predict completeNH_4_–N removal at temperatures above 20 °C while at 10°C nitrification appears as kinetically limited resulting in significant remaining NH_4_–N in the effluent despite the availability of DO.

In order to investigate the possible temperature effect on the treatment capacity of the wetland, different hydraulic loading rates were simulated. Figure 1.a shows the CR and CS predicted effluent concentrations at the different loadings rates and temperatures. CS concentrations are predicted to decrease with increasing temperature, as expected due to the faster hydrolysis rates at higher temperatures.

The predicted remaining CR increased with temperature and, to some extent, with loading rate however at loading rates between 2m^2^/PE and 4m^2^/PE these differences are minor. It is only after what could be described as the capacity limit threshold (corresponding to a point of loading rate between 2-1 m^2^/PE) when this effect becomes significant, the detailed spatial dynamic simulation results allowed us to attribute this behaviour to oxygen limitation as discussed below.

In regards to NH_4_-N removal, almost complete nitrification is achieved at the low loading rates 4m^2^/PE and 2m^2^/PE for all temperatures (Figure 1.b) with the expected positive kinetic effect of higher temperatures. The high loading rate of 1m^2^/PE simulations shows again the wetland treatment capacity limits being reached.

At low temperature (10°C) and despite of the higher oxygen solubility, a clear kinetic limitation of the microbial nitrification due to low growth rate and activity is predicted by the model, in consistency with literature (Akratos and Tsihrintzis, 2007). This limitation attenuates at higher temperature with better nitrification performance at 20°C (Figure 1.b). It is however the detailed modelling of the system, which allows for the prediction of a decrease on nitrification performance at the highest temperature (30°C) due to oxygen limitation analogous to that observed in COD removal.

The detailed HYDRUS CW2D model outputs allow for the investigation of the possible mechanisms behind the effects predicted above. Figure 2 shows the concentration of NH_4_–N and DO for 1 m^2^/PE loading rate at different temperatures in observation points at four different depths throughout the loading cycles. The loading cycle takes 15 minutes with a default time between loadings of 3 hours and starts right at the beginning of the data shown.

**Figure 2.**
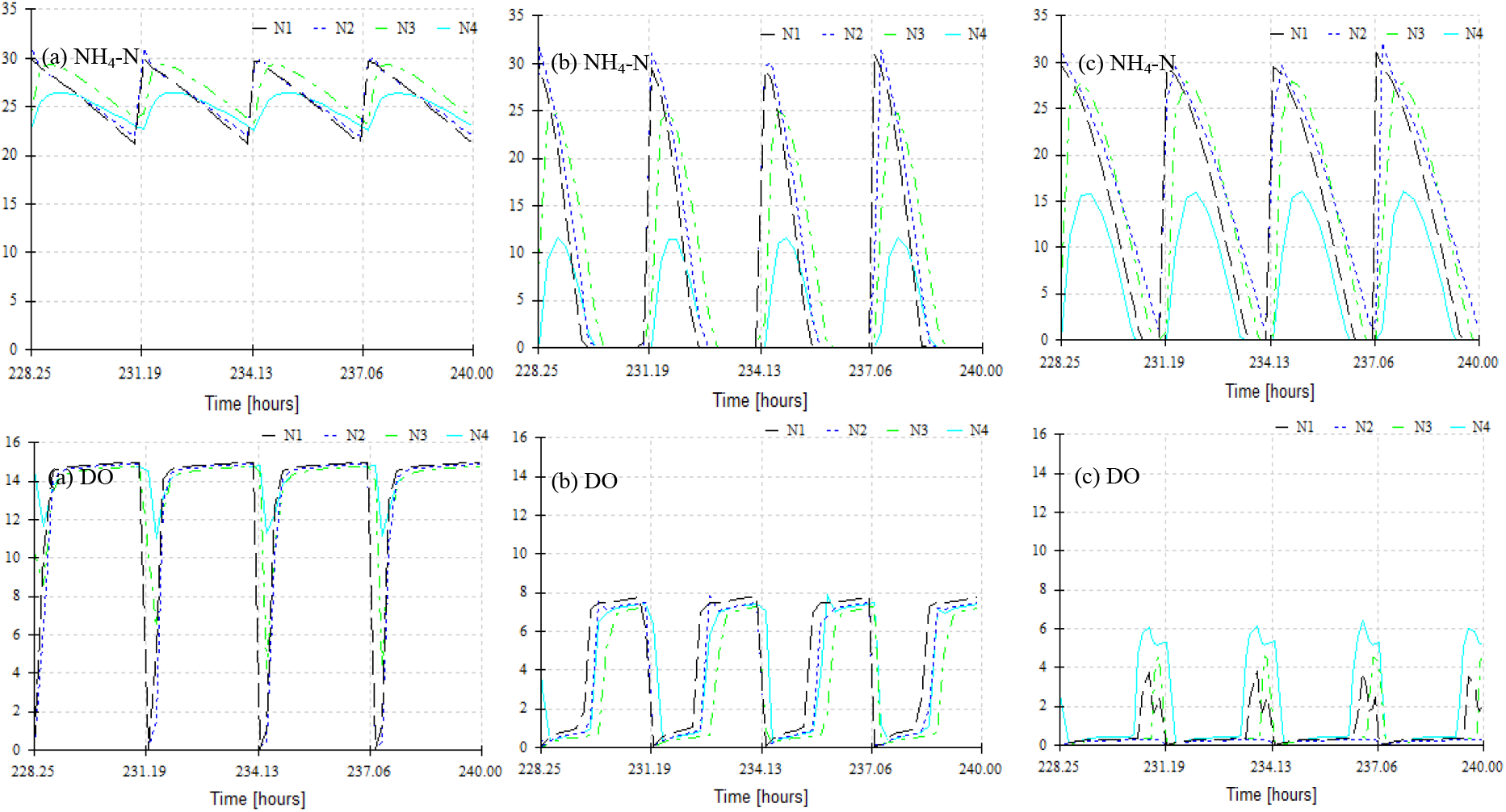
Simulated NH_4_–N and DO concentration profiles at different depths (observation points N1 = 4.96cm, N2 = 13.36cm, N3 = 25.92cm and N4 = 35cm) at (a) 10°C (b) 20 °C (c) 30 °C at a specific loading rate of 1 m^2^/PE.

Results at 10 °C (Figure 2(a)) show how NH_4_–N consumption is incomplete between loading cycles despite the high DO; this indicates a clear kinetic limitation of nitrifiers activity and growth. Results at 20 °C (Figure 2(b)) show that some NH_4_-N reaches the effluent during the first part of the loading cycle due to possible DO limitation while during the second part of the cycle DO is not limiting but no NH_4_ – N remains in the system. Results at 30 °C (Figure 2(c)) illustrate clearly the DO limitation throughout the whole cycle with no dead times in this case.

The simulation results indicate that nitrification in VSSFCW might be limited by DO during treatment at high temperatures. Results also show that possible dead times during cycles might reduce capacity suggesting that changes in the operation could optimise treatment capacity. The impact of loading frequency on the treatment capacity was evaluated through simulations at the maximum wetland capacity i.e. high loading rate (1m^2^/PE) at for the three different temperatures.

Despite literature suggesting no major correlation between temperature and COD removal (Akratos and Tsihrintzis, 2007; Chang *et al*., 2012), these simulation study suggest that temperature might indeed have an observable impact when the wetlands operate at their maximum capacity. These results suggest that a detailed experimental investigation of the effect of temperature should be conducted for CWs application in warm arid regions.

### Effect of loading frequency on treatment performance at maximum loading rate

A loading frequency of 8 day^−1^ with a 15 minute feeding time was used as default in the above simulations. For the study of the loading frequency impact on treatment performance three additional loading frequencies were simulated (4, 12 and 24 day^−1^) with adjusted feeding times to maintain the same daily loading rate to the wetland.

The simulation results predicted NH_4_–N effluent concentrations are higher at 30 °C than at 20 °C (Figure 1(b)) consequence of lower saturation concentration of oxygen. More frequent and less volume per loading is proposed to potentially increase nitrification capacity due to better availability of oxygen from the air entering with every loading. This was tested in simulation and results presented in Figure 3 in terms of NH_4_-N and DO. As the frequency of loading is increased, simulations predict lower DO concentrations at the higher layers (partly due to the shorter times between loadings not being sufficient for the water DO levels to reach the saturation at the higher layers) but an average higher DO in the lower layers. Simulations also predict average lower effluent NH_4_–N concentrations at higher loading frequencies possibly thanks to the average increased availability of DO in the lower layers. Similar (not shown) improved COD removal was predicted with increased loading frequency.

**Figure 3.**
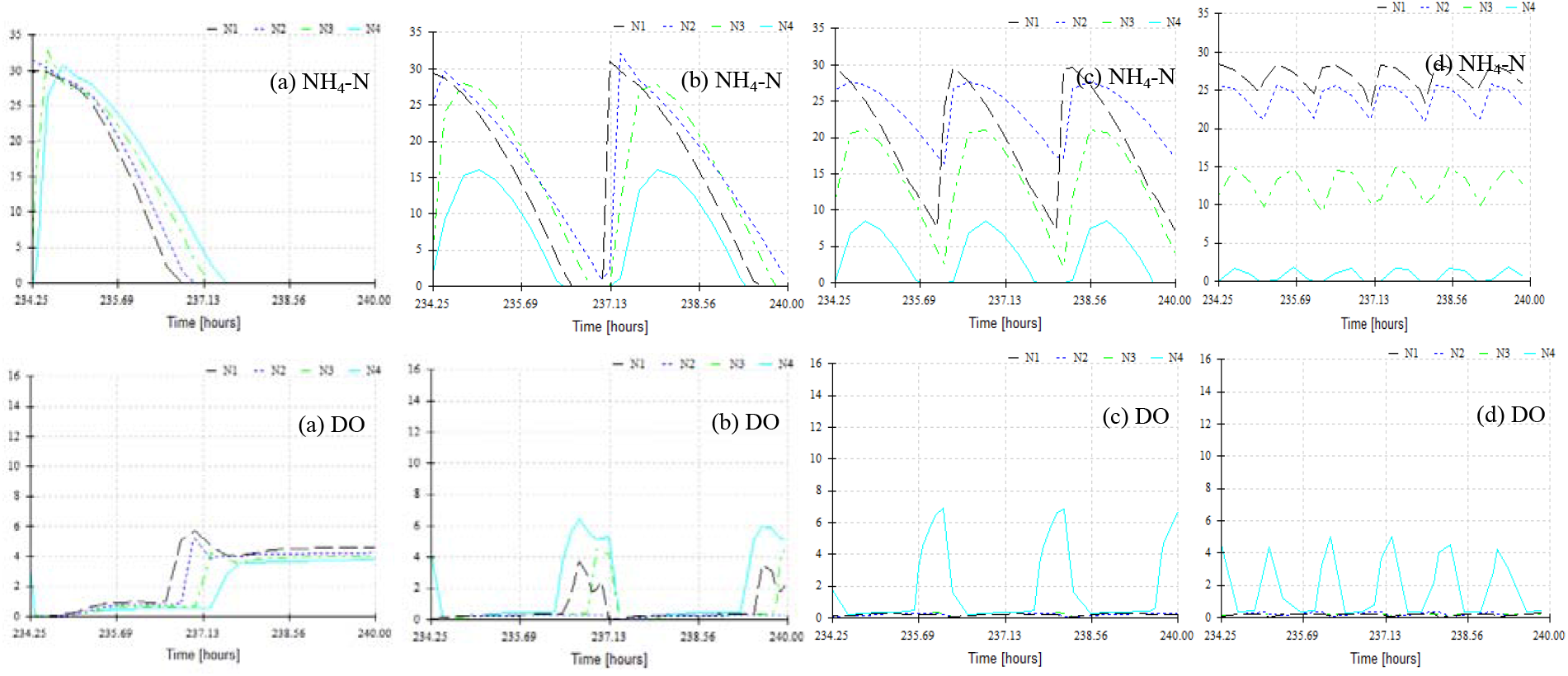
Simulated NH_4_ – N and DO concentration profiles at different depths (observation points N1 = 4.96cm, N2 = 13.36cm, N3 = 25.92cm and N4 = 35cm) at 30 °C and 1 m^2^/PE at frequencies of (a) 4 day^−1^ (b) 10 day^−1^ (c) 12 day^−^ 1 (d) 24 day^−1^

At low temperature (10 °C) no clear impact on the effluent concentration of NH_4_–N is predicted with only a slight decrease at frequencies both lower or higher than 8 day^−1^ (see Supplementary Figure S2). This is expected due to the temperature limited rate of nitrification at low temperatures.

At moderate temperature (20°C) lower simulated NH_4_–N effluent concentrations are reached at higher frequency of loading. Results presented in Supplementary Figure S3 show frequencies higher of 8, 10 and 12 day^−1^ to provide sufficient time between two consecutive loadings appears as sufficient to achieve almost complete nitrification while simultaneously avoiding dead times with no activity in between cycles. Low frequencies appear to lead to most of the water flowing out without NH_4_–N removal and a reduction in the overall average effluent quality.

All these model-based results suggest that an optimum operation exists in terms of loading frequency that would need to be evaluated experimentally for each specific wetland to achieve, particularly at high temperature, an overall improved treatment performance. The model proved of potential value to describe interactions between flows, NH_4_-N and air supply to the system.

The above combined simulations results provide hypotheses and guidelines for the investigation of several non-trivial interactions in VSSFCWs, potentially useful to optimise the design and operation for warm arid regions. In these regions CWs operate at temperatures much higher than those from which most existing guidelines were developed.

## Conclusion

The simulation study using HYDRUS CW2D provides a mechanistic insight into the possible effects of temperature, loading rate and loading frequency on the performance of VSSFCWs in terms of COD removal and NH_4_-N. These insights can be of use to formulate hypotheses and operational strategies to improve the overall treatment performance and capacity of VSSFCW, particularly in warm climates.

Simulations suggest that the impact of temperature on both COD and NH_4_-N removals may occur through two mechanisms, namely low growth microbial kinetic limitations at low temperatures and DO concentration limitations at high temperatures. The detailed spatial simulation of the system allowed also for the identification of possible dead times causing lower performance if low loading frequencies are used. The modelling study suggests that specific experiments should be conducted in each wetland to determine the optimum loading frequencies in order to adjust treatment capacity and performance through improved oxygen transfer, (especially at high temperatures) and minimum dead times.

## Acknowledgements

The Sustainable Bioenergy Research Consortium (SBRC) and the Government of Abu Dhabi for the project funding. Drs Jirka Šimunek and Günter Langergraber for their support on questions related to HYDRUS and the wetland module.

## Notes

### Competing Interest Statement

The authors have declared no competing interest.

