## Supplementary material for "Effect of temperature and loading frequency on the performance of vertical subsurface flow constructed wetlands: Modelling study using HYDRUS": Suplementary Information

**Table S1**. Kinetic Parameters in CW2D (Langergraber & Šimůnek, 2005)

| Kinetic parameter | Description | 10°C | 20°C | 30°C | |
|---|---|---|---|---|---|
| | | Literature | Literature | Estimated | Used |
| $K_h$ [1/d] | hydrolysis rate constant | 2.00 | 3.00 | 4.38 | 4.38 |
| $K_X$ [$mg_{COD,CS}/mg_{COD,BM}$] | saturation/inhibition coefficient for hydrolysis | 0.22 | 0.10 | 0.05 | 0.05 |
| $\mu_H$ [1/d] | maximum aerobic growth rate on CR | 3.00 | 6.00 | 11.46 | 11.46 |
| $b_H$ [1/d] | rate constant for lysis | 0.20 | 0.40 | 0.76 | 0.76 |
| $K_{het,O2}$ [$mg_{O2}$/l] | saturation/inhibition coefficient for $O_2$ | 0.20 | 0.20 | - | 0.20 |
| $K_{het,CR}$ [$mg_{COD,CR}$/l] | saturation/inhibition coefficient for substrate | 4.00 | 4.00 | - | 4.00 |
| $K_{het,NH4N}$ [$mg_{NH4N}$/l] | saturation/inhibition coefficient for NH4 (nutrient) | 0.05 | 0.05 | - | 0.05 |
| $K_{het,IP}$ [$mg_{IP}$/l] | saturation/inhibition coefficient for P | 0.01 | 0.01 | - | 0.01 |
| $\mu_{DN}$ [1/d] | maximum denitrification rate | 2.40 | 4.80 | 9.17 | 9.17 |
| $K_{DN,O2}$ [$mg_{O2}$/l] | saturation/inhibition coefficient for $O_2$ | 0.20 | 0.20 | - | 0.20 |
| $K_{DN,NO3N}$ [$mg_{NO3N}$/l] | saturation/inhibition coefficient for $NO_3$ | 0.50 | 0.50 | - | 0.50 |
| $K_{DN,NO2N}$ [$mg_{NO2N}$/l] | saturation/inhibition coefficient for NO2 | 0.50 | 0.50 | - | 0.50 |
| $K_{DN,CR}$ [$mg_{COD,CR}$/l] | saturation/inhibition coefficient for substrate | 4.00 | 4.00 | - | 4.00 |
| $K_{DN,NH4N}$ [$mg_{NH4N}$/l] | saturation/inhibition coefficient for NH4 (nutrient) | 0.05 | 0.05 | - | 0.05 |
| $K_{DN,IP}$ [$mg_{IP}$/l] | saturation/inhibition coefficient for P | 0.01 | 0.01 | - | 0.01 |
| $\mu_{ANs}$* [1/d] | maximum aerobic growth rate on $NH_4N$ | 0.30 | 0.90 | 2.51 | 2.51 |
| $b_{ANs}$* [1/d] | rate constant for lysis | 0.05 | 0.15 | 0.42 | 0.42 |
| $K_{ANs,O2}$* [$mgO_2$/l] | saturation/inhibition coefficient for O2 | 1.00 | 1.00 | - | 1.00 |
| $K_{ANs,NH4N}$** [$mg_{NH4N}$/l] | saturation/inhibition coefficient for $NH_4$ | 5.00 | 0.50 | - | 0.50 |
| $K_{ANs,IP}$ [$mg_{IP}$/l] | saturation/inhibition coefficient for P | 0.01 | 0.01 | - | 0.01 |
| $\mu_{ANb}$* [1/d] | maximum aerobic growth rate on $NO_2N$ | 0.35 | 1.00 | 2.67 | 2.67 |
| $b_{ANb}$* [1/d] | rate constant for lysis | 0.05 | 0.15 | 0.42 | 0.42 |
| $K_{ANb,O2}$* [$mg_{O2}$/l] | saturation/inhibition coefficient for $O_2$ | 0.10 | 0.10 | - | 0.10 |
| $K_{ANb,NO2N}$* [$mg_{NO2N}$/l] | saturation/inhibition coefficient for $NO_2$ | 0.10 | 0.10 | - | 0.10 |
| $K_{ANb,NH4N}$ [$mg_{NH4N}$/l] | saturation/inhibition coefficient for $NH_4$ (nutrient) | 0.05 | 0.05 | - | 0.05 |
| $K_{ANb,IP}$ [$mg_{IP}$/l] | saturation/inhibition coefficient for P | 0.01 | 0.01 | - | 0.01 |

**Table S2**. Wetland Characteristics

| Variable | CW |
|---|---|
| Dimensions | 1m (length) x 1m (width) x 0.5m (height) |
| Hydraulic Loading Rate | 200 L/m$^2$ – day or 0.2 m$^3$/m$^2$ – day |
| Number of Loadings | 8 day $^{-1}$ |
| Loading time | 15 minutes |
| Filter Layer | Double washed wadi sand (gravel size 0 – 4 mm) 35 cm deep |
| Drainage layer | Gravel (size 5 – 10 mm) 15 - 30 cm deep |

**Table S3**. Soil Hydraulic Parameters of the van Genuchten – Mualem model (Langergraber & Šimůnek, 2005)

| Parameter | Residual Water Content $\theta_r$ (m$^3$ / m$^3$) | Saturated Water Content $\theta_s$ (m$^3$ / m$^3$) | Shape Parameters α (m$^{-1}$) | n | l | Saturated hydraulic conductivity $K_s$ (m/h) |
|---|---|---|---|---|---|---|
| Measured $K_s$ + porosity | 0.045 | 0.3 | 14.5 | 2.68 | 0.5 | 1.17 |

Table S4. Evaporation and Soil Minimum Pressure

| Parameter | Unit | Symbol | Values | | |
|---|---|---|---|---|---|
| Temperature | °C | T | 10 | 20 | 30 |
| Evaporation rate | cm/h | E | 1.62E-02 | 1.71E-02 | 1.18E-01 |

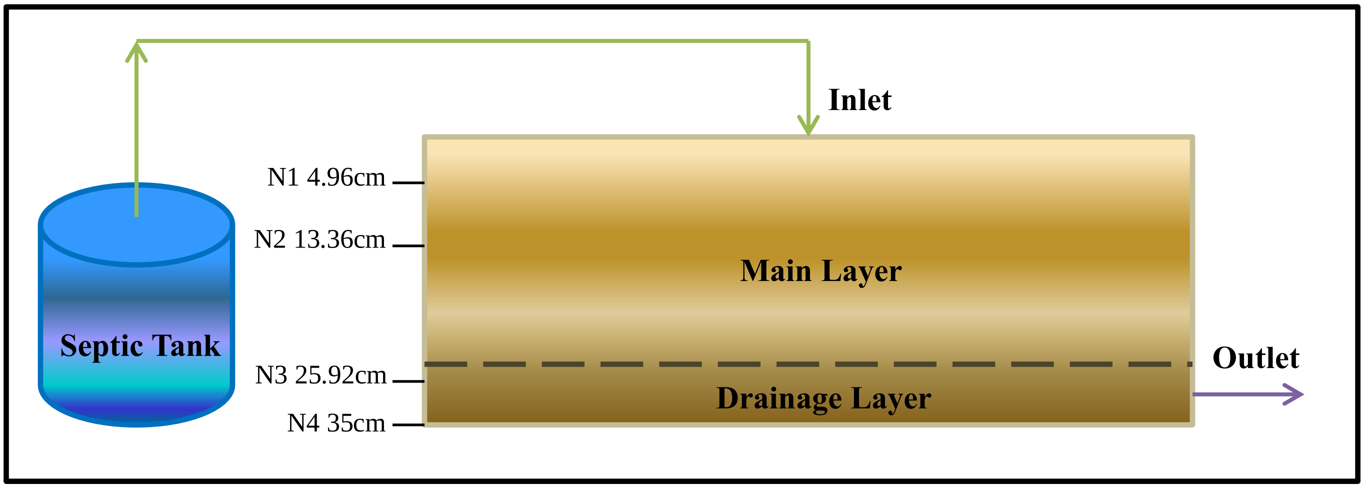

**Figure S1**. Modelled vertical subsurface flow constructed wetland treating septic tank effluent. Observation nodes at different depths are indicated.

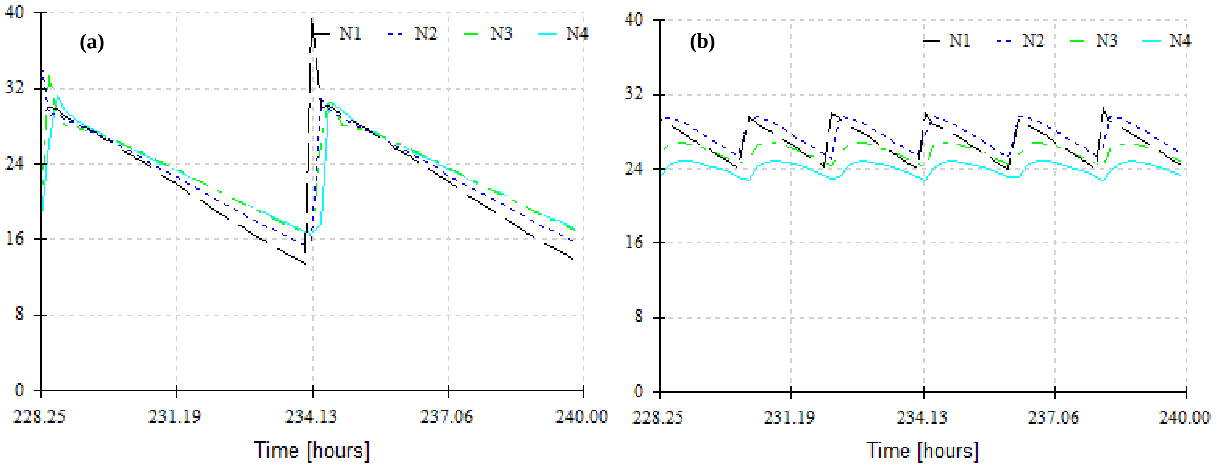

**Figure S2**. $NH_4$–N concentration profiles at different depths (observation points N1 = 4.96cm, N2 = 13.36cm, N3 = 25.92cm and N4 = 35cm) at 10 °C and 1 $m^2$/PE at frequencies of (a) 4 $day^{-1}$ (b) 12 $day^{-1}$

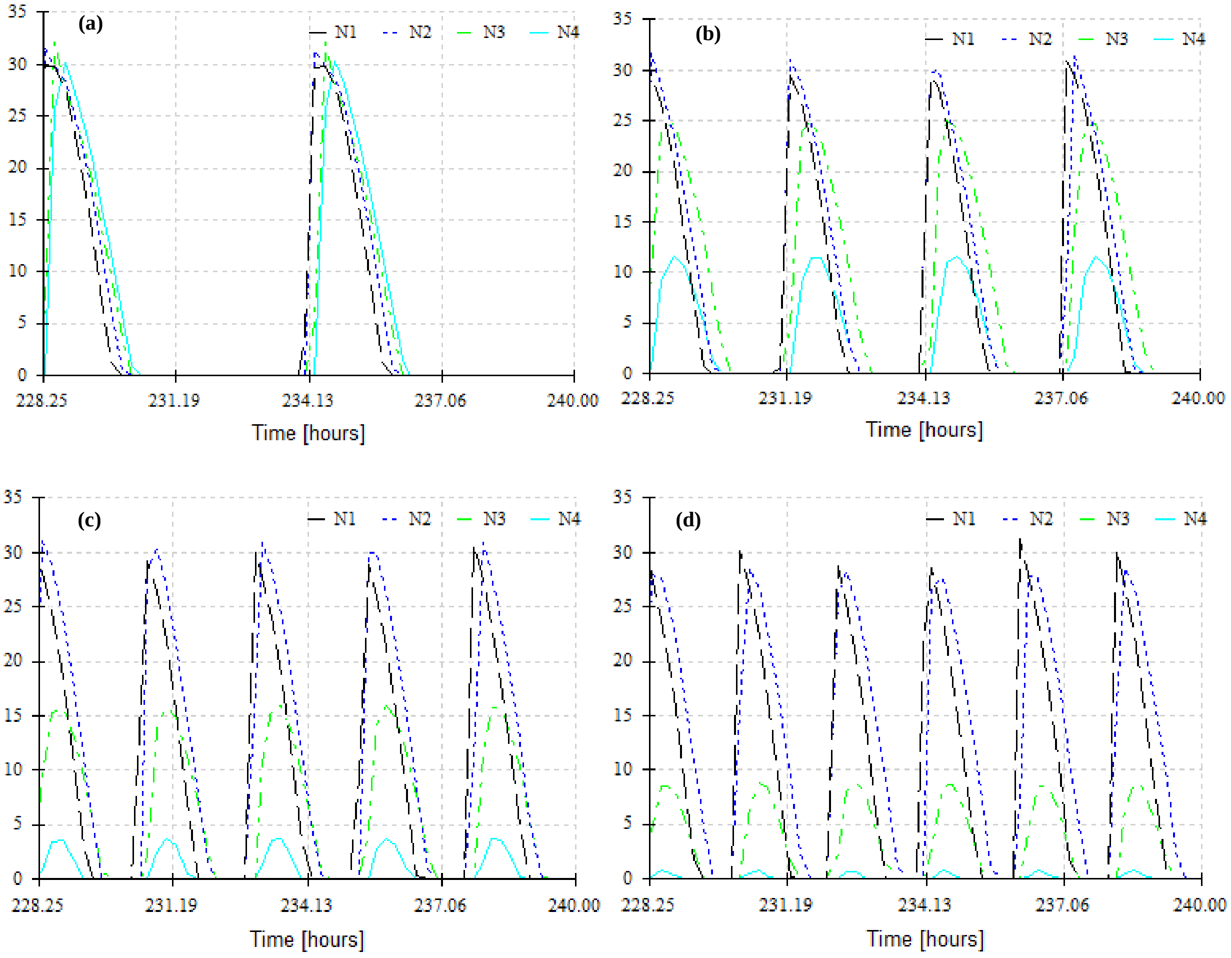

**Figure S3**. $NH_4$–N concentration profiles at different depths (observation points N1 = 4.96cm, N2 = 13.36cm, N3 = 25.92cm and N4 = 35cm) at 20 °C and 1 $m^2$/PE at frequencies of (a) 4 day-1 (b) 8 day-1 (c) 10 day-1 (d) 12 day-1

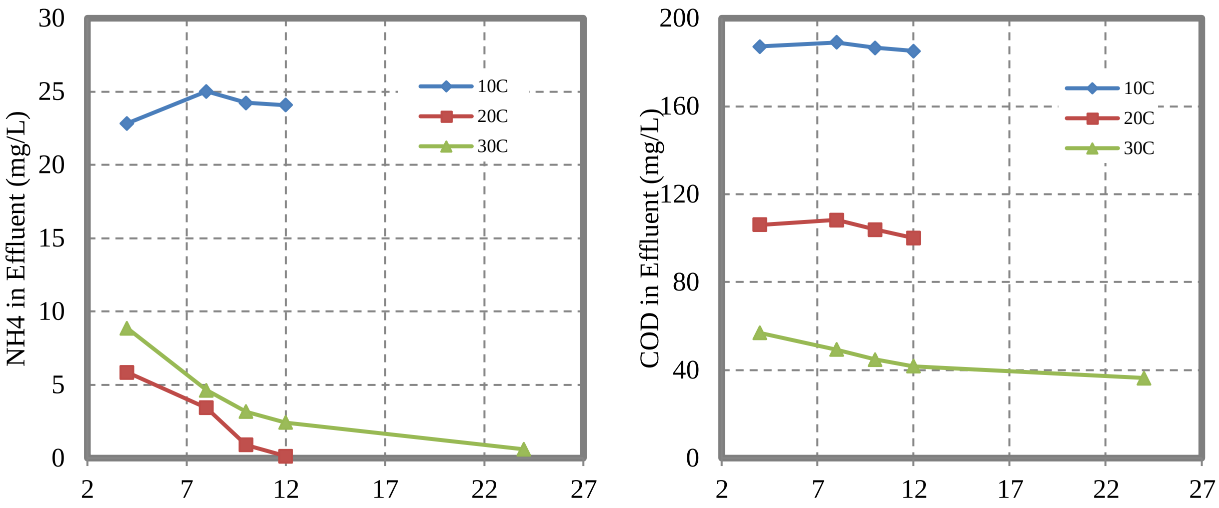

**Figure S4**. Simulated effluent concentrations (a) $NH_4$–N and (b) COD at a specific loading rate of 1 $m^2$/PE at different loading frequencies and temperatures.
